# Ancient secretory pathways contributed to the evolutionary origin of an ecologically impactful bioluminescence system

**DOI:** 10.1101/2024.05.07.593075

**Authors:** Lisa Y. Mesrop, Geetanjali Minsky, Michael S. Drummond, Jessica A. Goodheart, Stephen R. Proulx, Todd H. Oakley

**Affiliations:** Department of Ecology, Evolution, and Marine Biology, University of California Santa Barbara, CA, 93106 U.S.A; Division of Invertebrate Zoology, American Museum of Natural History, New York, NY, 10024 USA

**Keywords:** novelty, evolution, bioluminescence, WGCNA, complex traits, evo-devo

## Abstract

Evolutionary innovations in chemical secretion – such as the production of secondary metabolites, pheromones, and toxins – profoundly impact ecological interactions across a broad diversity of life. These secretory innovations may involve a “legacy-plus-innovation” mode of evolution, whereby new biochemical pathways are integrated with conserved secretory processes to create novel products. Among secretory innovations, bioluminescence is important because it evolved convergently many times to influence predator-prey interactions, while often producing courtship signals linked to increased rates of speciation. However, whether or not deeply conserved secretory genes are used in secretory bioluminescence remains unexplored. Here, we show that in the ostracod *Vargula tsujii*, the evolutionary novel c-luciferase gene is co-expressed with many conserved genes, including those related to toxin production and high-output protein secretion. Our results demonstrate that the legacy-plus-innovation mode of secretory evolution, previously applied to sensory modalities of olfaction, gustation, and nociception, also encompasses light-producing signals generated by bioluminescent secretions. This extension broadens the paradigm of secretory diversification to include not only chemical signals but also bioluminescent light as an important medium of ecological interaction and evolutionary innovation.

**Significance Statement:** Animals produce an enormous diversity of secreted chemical products, like toxins and pheromones, with wide-ranging impacts on ecological interactions. Although a deeply conserved toolkit of secretory genes may often underlie chemical interactions mediated through smell, taste, and sensing pain, whether or not this evolutionary mode generalizes to sensing light is unknown. Here we show that a bioluminescence secretion system, which creates light for anti-predator and courtship interactions, also uses genes of a deeply conserved secretory toolkit. Therefore, secretory innovations may act through all sensory modalities by integrating conserved genes with novel biosynthesis pathways, to serve as crucibles of evolutionary and ecological diversity.

## Introduction

Exocrine glands are specialized secretory structures, often representing lineage-specific innovations that produce diverse chemical secretions. These secretions – including secondary metabolites, pheromones, and toxins – have been instrumental in evolving new ecological interactions using behaviors like intraspecific communication (e.g. courtship signaling and caste social systems) and predator-prey interactions (1–3). Examples of secretory innovations include courtship glands of salamanders, venom glands of snakes and remipedes, defensive tergal glands of rove beetles, and ink and opaline glands of mollusks (4–8). Secretory innovations may often use a shared “genetic toolkit” (9) for secretion that was modified in different lineages by adding new genes to produce diverse products (10), a mode of evolution we call “legacy-plus-innovation”. Although exocrine glands clearly exhibit widespread morphological convergence, only a few studies have characterized gene expression to support the use of a shared genetic toolkit and the legacy-plus-innovation model (6, 8, 10), limiting our understanding of the historical constraints that shape the broad range of secretory novelties.

Thus far, the legacy-plus-innovation mode of secretory evolution is proposed for secretory outputs perceived by olfactory, gustatory, and nociceptive mechanisms (10), leaving the genetic underpinnings of light-producing secretions largely unexplored, despite the ecological importance of visual interactions (reviewed in 11). Chemical secretions that generate light are produced by bioluminescent systems, which have evolved convergently many times, using an impressive variety of structural and functional forms (12, 13). Bioluminescent glands facilitate many ecological interactions, including courtship signals, anti-predation ‘burglar alarm’ displays, and various predation strategies (3, 14–17). For example, bioluminescent glands of some marine crustaceans and syllid worms discharge glowing mucus into the water as anti-predation and courtship displays and the bioluminescent glands of pocket sharks secrete illuminated lures to attract prey (14, 15, 18). The widespread evolution of secretory bioluminescence and the ecological consequences associated with visual interactions of light displays motivate a quest to understand the genetic underpinnings of these adaptive and often beautiful secretions.

Secretory bioluminescence highlights vision as an important sensory modality within the framework of secretory evolution, but the extent to which a shared secretory toolkit also underlies secreted bioluminescence remains obscure. To test whether “legacy-plus-innovation” underlies secretory bioluminescence, we analyze gene expression in the bioluminescent upper lip of cypridinid ostracods (crustaceans), a secretory innovation with important ecological, cellular, and biochemical functions (19–21). Since its origin with the clade Luminini at least 237 MYA (22), luminine bioluminescence has had important ecological functions, including deterring predation and signaling for courtship. Nearly all Luminini release large amounts of bioluminescent mucus during predation attempts, often escaping unscathed, due either to the light itself and/or to the unpalatability of the mucus (15). Within Luminini, a subclade called Luxorina also produces complex bioluminescent signals for courtship, which only evolved once (22, 23). The bioluminescence of Luminini is produced by secretory cells and nozzle-like appendages embedded in the upper lip (labrum) (20, 21). While the upper lip as a whole is homologous between bioluminescent and non-bioluminescent ostracods, the luminous upper lip is a light organ unique to Luminini, and therefore may be considered a radical transformation of an existing body part, which Wagner (24) terms a Type-II novelty. Compared to non-bioluminescent upper lips, bioluminescent upper lips have more cell types, including those that synthesize and secrete mucus that contains light-emitting compounds (20, 21). The two compounds include evolutionarily novel (25), biochemically well-characterized enzymes called c-luciferases (26–28); and a novel small molecule that is oxidized to produce light, a luciferin often called vargulin (29).

In addition to the established roles of novel c-luciferases and vargulin in evolving secretory bioluminescence, we herein report that the upper lip of a bioluminescent ostracod expresses deeply conserved genes of secretory pathways, despite vast evolutionary distances separating this bioluminescence system from other secretory innovations. Using the bioluminescent ostracod *Vargula tsujii*, we analyzed genes co-expressed with c-luciferase, which we call a “Bioluminescent Co-Regulatory Network” (BCN). We find this BCN to contain many conserved and putatively secreted genes, some of which resemble toxin-like products; as well as non-secreted housekeeping genes involved in pathways related to protein secretion and associated stress responses. We also compared differential gene expression of entire upper lips of luminous and non-luminous ostracods, finding distinct patterns of differential expression in the luminous upper lip. The differential expression analyses further emphasize the use of conserved toxin-like genes and other conserved secretory pathways. At the same time, we report many novel genes besides c-luciferase are expressed in the BCN and luminous upper lip. Together, these results reveal legacy-plus-innovation evolution in the bioluminescent upper lip, whereby elements of a deeply conserved secretory toolkit are deployed along with new genes within new co-expression networks. Our results extend this secretory innovation paradigm (10) to also encompass secreted light as a crucial medium of ecological interaction and evolutionary innovation.

## Results

### Co-expression of conserved secretory pathways and toxin-like genes with c-luciferase

The genes co-expressed with c-luciferase across 51 tissues/stages of *V. tsujii* form a co-expression network module with 890 genes that we call the “Bioluminescent Co-Expression Network’’ (BCN), of which 26% exhibit high module membership (MM>.8) (**Fig. 1A, Dataset S1 A**,**D**). The network revealed strong connections (MM>.8) between c-luciferase, luciferase-like genes, toxin-like genes, other putatively secreted products (based on presence of a signal peptide), and non-secreted housekeeping genes **(Fig. 1A)**. In addition to bioluminescence, significantly enriched GO terms of the BCN include protein transport and modification, lipid biosynthesis, peptide modifications, tissue development, cell signaling, hydrogen peroxide breakdown, response to stimuli and cellular stress, and oxidative stress and inflammation **(Fig. 1B, Dataset S1 F-H**). Annotations of non-secreted housekeeping genes include maintenance of tissue and cellular functions, involvement in protein secretory pathways that respond to cellular stress, and modulation of stress response pathways (**Fig. 1B)**. The concerted activity of these processes could be related to demands of protein synthesis and secretion consistent with the role of the bioluminescent upper lip in high-volume secretion of c-luciferase and other proteins integral to the bioluminescent mucus. Other biological processes enriched in the BCN include neuromuscular synaptic transmission **(Fig. 1B)**, perhaps from labral nerves that innervate the upper lip (30). We found no evidence for the conservation of co-expression of BCN orthologs compared to similar networks expressed in the non-luminous relative (***S1 Appendix, Fig*.*S1***).

**Figure 1.**
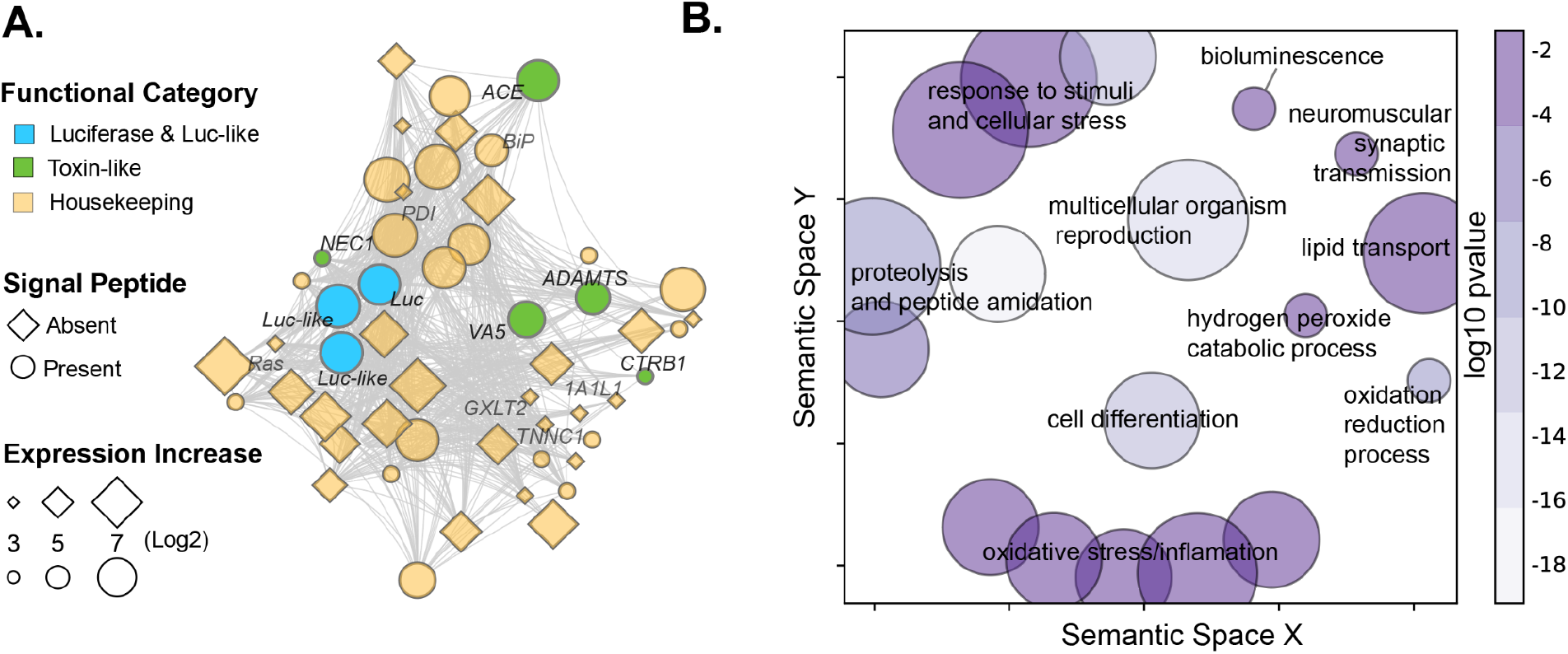
**(A)** The bioluminescent co-regulatory network (BCN) represents a group of co-expressed genes that are associated with the expression of luciferase, luciferase-like genes, toxin-like genes, and housekeeping genes some of which are constituents of stress and secretory protein pathways. The BCN comprises over 800 co-expressed genes and more than one-third of the BCN are significantly upregulated uniquely in the upper lip of *V. tsujii* and are considered integral to the network (MM > 0.8). Out of 227 genes uniquely upregulated in the bioluminescent upper lip, only genes that have annotations (26.4 %, 60 annotated genes) were plotted to visualize connections and overall network topology. Bioluminescent genes, including c-luciferase (Luc) and phylogenetically similar but not functionally tested luciferase-like (Luc-like) genes are indicated by the color blue. Toxin-like genes include Venom Allergen 5 (VA5), A disintegrin and metalloproteinase with thrombospondin (ADAMTS), Chymotrypsinogen B (CTRB1), Neuroendocrine convertase 1 (NEC1), Angiotensin-converting enzyme (ACE) is indicated by the color green. Genes in the protein secretory pathway include Protein disulfide-isomerase 2 (PDI), heat shock protein 70 (Hsp70; BiP), Ras and EF-hand domain-containing protein (RAS), Glucoside xylosyltransferase 2 (GXLT2), Troponin C (TNNC1),1-aminocyclopropane-1-carboxylate synthase-like protein 1 (1A1L1) are labeled in yellow. Genes that have a signal peptide and are presumably secreted outside the cell are indicated by a circle and genes that do not have a signal peptide are indicated by a diamond. Distance between nodes indicates strength connectivity. **(B)** GO terms enriched in the BCN are visualized using GO-Figure. Each bubble represents a GO term or a cluster of GO terms summarized by a representative term reported in the supplement. The size of the bubble indicates the number of GO terms in each cluster and the color is the P value or the average P value of the representative GO term in that cluster. The GO terms for each bubble are found in **Dataset S1 H**.

Our analyses reveal for the first time the co-expression of c-luciferase with deeply conserved toxin-like domains and genes, which is of particular interest because ostracods use bioluminescence in anti-predator responses. In *V. tsujii*, we find the BCN to have transcripts with signal peptides similar to genes expressed in venom glands and salivary glands of other animals. Out of the 105 BCN transcripts (**Dataset S1 B**) with a signal peptide and no transmembrane domain (and therefore likely to be secreted products) 23 have domains of known toxin gene families (7) (**Dataset S2 B, K**). At least 10 (37% of known toxin gene families) of these toxin-like genes have high connectivity (MM > 0.8) in the BCN (i.e., hub genes), with several also significantly upregulated in the bioluminescent upper lip (**Dataset S1 D** and **Dataset S2 K**). Like the ‘metavenom network’ expressed in the venom glands of snakes (8), we find many genes in the BCN with similarity to genes involved in the global endoplasmic reticulum (ER) stress (ERS) response, which is also part of the protein secretory pathway. Genes of the Unfolded Response Pathway (UPR) and ER-associated degradation (ERAD) pathways that are responsible for high-output protein secretion also have strong connections with c-luciferase and luciferase-like genes in the BCN: We found transmembrane emp24 domain (TMED), endoplasmic reticulum chaperone BiP (HSP70), protein disulfide isomerases (PDI), translation initiation factor 2 (IF2) and ubiquitin ligases **(Fig. 1A, Dataset S3 A, B**). We also identified deeply conserved genes associated with processes downstream of the ERS response pathways such as detoxification, including oxidation-reduction genes such as aldehyde dehydrogenase, sulfotransferases, cytochrome P450s, hydroxylases, and conjugation and hydrolytic enzymes (**Fig. 1B, Dataset S1 A, F-H**) (31, 32). One sulfotransferase (Vt-LST3) that sulfates vargulin *in vitro* (33) is co-regulated with c-luciferase in the BCN and is upregulated in the bioluminescent upper lip compared to the eye and gut, is the best candidate for creating a storage form of vargulin in *V. tsujii* (**Dataset S1 A, Dataset S4 A)**.

### Differential expression of conserved secretory pathway genes and toxin-like gene families

The results of expression analyses in upper lips of a luminous *V. tsujii* and non-luminous *Skogsbergia sp*. reveal an intriguing mix of legacy and innovation. The common evolutionary legacy of upper lips is reflected in the similar overall expression patterns of one-to-one orthologs between the upper lips of the two species, which cluster by tissues **(Fig. 2A)**. In addition, both species show significantly upregulated genes that comprise an unexpectedly complex cocktail of transcripts coding for putative toxin-like proteins with signal peptides. For the bioluminescent upper lip, out of the total number of upregulated transcripts with signal peptides and without transmembrane domains, we found 25 transcripts (26%) representing at least 11 different toxin-like classes **(Fig. 3A, Dataset S2 B, C, E)**. The 11 different toxin classes can be classified broadly into three functional categories: 1) neurotoxins: ShKT; 2) protease inhibitors: Kunitz, lipocalin, 3) Other enzymes: peptidase S1, peptidase S8, peptidase M2, metallopeptidase M12, CUB, Serpin and C-type Lectin and 4) CAP domain proteins. For the non-bioluminescent upper lip, we found 13 transcripts (15.3%), including some from classes shared with the bioluminescent upper lip: C-type Lectin, ShKT, CUB, WAP, Peptidase S1, Peptidase S8, and metallopeptidase M12 **(Fig. 3B, Dataset S2 B, D, H)**. For both species, we compared the ratio of putative toxin-like genes to secreted genes in both upper lips compared to two organs not involved in the secretory process of anti-predator displays -- the compound eye and gut. For the luminous *V*.*tsujii*, we found the ratio of putative toxin-like genes to secreted genes to be significantly higher in the bioluminescent upper lip compared to the gut, but not significantly higher in the compound eye (BUL:Compound Eye; Fisher’s Exact Test; p=0.1371, BUL:Gut; p<=0.035)(**Fig. 3A, Dataset S2 L**). In contrast, the non-luminous upper lip expressed a lower ratio of putative toxin-like genes to secreted genes compared to the compound eye and gut (**Fig. 3B**)

**Figure 2:**
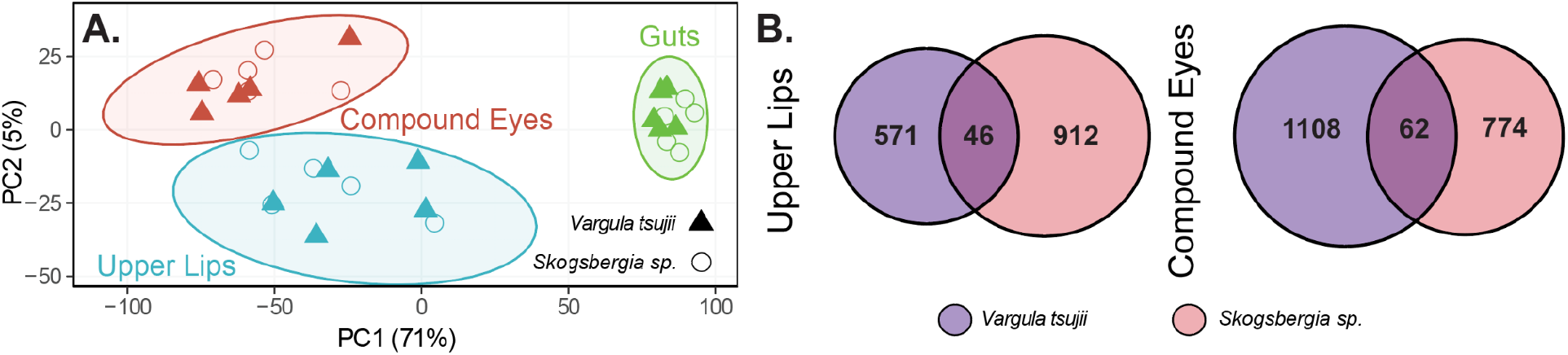
Orthologous gene expression values from two species cluster by tissues, while differentially expressed genes mainly differ between upper lips of two species **(A)** Expression patterns of orthologous genes between upper lips of bioluminescent and non-bioluminescent ostracods are conserved. PCA clustered gene expression of 4217 one-to-one orthologs by tissue, and differences among tissues explain more than 75 % of the variation in the dataset. Ellipses represent 95% confidence intervals. **(B)** Venn diagram illustrating the number of uniquely significantly upregulated genes shared across both upper lips vs compound eyes for each species, *Vargula tsujii* and *Skogsbergia sp*.

**Figure 3:**
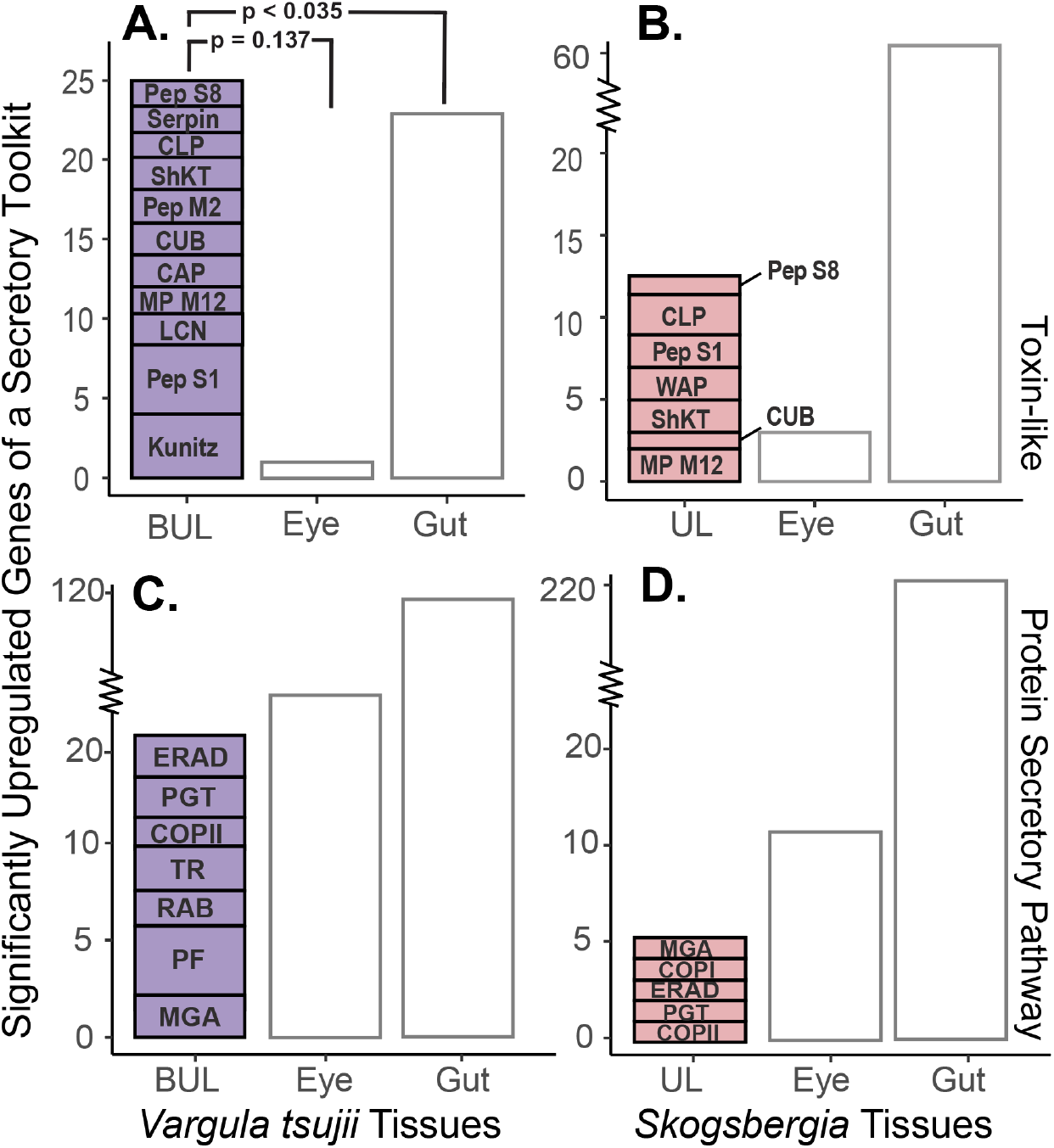
Differential expression of toxin-like secretory and secretory protein pathway products across luminous and non-luminous upper lips. **(A)** Compared to the gut, the luminous upper lip has a significantly higher proportion of toxin-like genes to secreted genes, but not significantly higher compared to the compound eye. Statistically significant pairwise comparison between BUL:Gut is indicated by p<=0.035. The luminous upper lip has at least 25 upregulated genes that have domains that belong to known toxin gene families which include Peptidase S8 (Pep S8), Serpin, C-Type Lectin (CLP), ShKT, Peptidase M2 (Pep M2), CUB, CAP, Metallopeptidase M12 (MP M12), Lipocalin (LCN), Peptidase S1 (Pep S1), Kunitz. **(B)** Compared to the compound eye and gut, the non-luminous upper lip expressed a lower proportion of toxin-like genes to secreted genes. The non-luminous upper lip has at least 13 upregulated genes that belong to known toxin gene families which include Peptidase S1 (Pep S8), C-Type Lectin (CLP), Peptidase S1 (Pep S1), WAP, ShKT, CUB and Metallopeptidase M12 (MP M12)**(C)** The luminous upper lip has at least 20 upregulated genes that belong to 7 different functional modules that belong to the secretory protein pathway which include: Golgi glycosylation (MGA), trafficking regulation (RAB), protein folding (PDI), post-Golgi trafficking (PGT), ERAD, coat protein complex I (COPI), and translocation (TR). **(D)** The non-luminous upper lip only has at least 5 genes that belong to the protein secretory pathway modules MGA, COPI, ERAD, PGT, and COPII. *BUL - Upper Lip*.

Although sharing similar overall expression of orthologs and general classes of expressed toxin-like genes and some semantically similar GO terms (**Dataset S4 K, *S1 Appendix, Fig*.*S2-S4***), we found only 46 specific genes to be shared between both upper lips in comparisons of all significantly upregulated genes uniquely expressed, compared to 62 shared between the eyes of the same two species (Fisher’s Exact Test; p-value = 0.8441) **(Fig. 2B, Dataset S4 J)**. Semantically similar GO terms across both upper lips include processes related to proteolysis, protein hydroxylation, tissue development and signaling, and metabolic and neural-related processes (**Dataset S4 G, *S1 Appendix, Fig*.*S2-S4***). Among the genes shared across both upper lips, several are putatively secreted genes similar to toxin and anticoagulant proteins, with a few associated with proteolysis. A major difference in gene expression between the two upper lips is that the luminous upper lip contains a significantly higher number of upregulated genes of the deeply conserved protein secretory pathway (**Fig. 3C, 3D**). The protein secretory pathway can be compartmentalized into functional modules responsible for protein folding, post-translational modifications, and trafficking of proteins. In the bioluminescent upper lip, we found at least 20 genes that belong to 7 different functional modules including, trafficking regulation (RAB), protein folding, ERAD, post-Golgi trafficking (PGT), coat protein complex I (COPI), coat protein complex II (COPII) **(Fig. 3C, Dataset S3 A, C)**. The UPR and ERAD pathways induce multiple downstream stress response genes including antioxidant, detoxification, and apoptosis (32),(34). We found several antioxidant and detoxification genes, and genes related to the apoptotic pathway upregulated in the bioluminescent upper lip. Although there are more secretory protein pathway genes upregulated in *V. tsujii* upper lip compared to *Skogerbergia* upper lip, and the number of gene copies varied across a taxonomically wider set of bioluminescent and non-bioluminescent lineages, we did not observe a clear trend between gene copy number and bioluminescent lineages (**Fig. 3C, 3B, *S1 Appendix, Fig*.*S5***).

### The novel bioluminescent upper lip expresses a higher proportion of novel genes

In addition to ancient genes of a conserved secretory toolkit, the bioluminescent upper lip and the BCN express many novel genes that originated similarly in time to bioluminescence itself (e.g. Luminini-specific genes, **Fig. 4**). First, a high proportion of all genes upregulated in the *V. tsujii* upper lip are Luminini-specific. More precisely, the proportion of Luminini-specific genes upregulated in the bioluminescent upper lip is significantly higher than the proportion of novel genes upregulated in the compound eye or gut, two tissues that are much older than the origin of bioluminescence (BUL:Compound Eye, Fisher’s exact test, p<0.0001; BUL:Gut, Fisher’s exact test, p<0.0034) **(Fig. 4B, Dataset S5 B)**. Second, compared to the complete set of expressed genes (e.g. expressed in luminous upper lip, gut, and compound eye), only the upregulated genes of the bioluminescent upper lip were characterized by a significantly higher proportion of Luminini-specific genes (BUL:DGE Dataset, Chi-Square, p<0.0002) **(Fig.4B, Dataset S5 B)**. Finally, both the upper lip and the BCN contain a high number of Luminini-specific genes. More precisely, the total number of Luminini-specific genes in the BCN is numerically higher than novel genes from a co-expression module of the gut, although not significant (BCN:Gut Module, Fisher’s Exact Test; p = 0.58) **(Fig. 4C, Dataset S5 C)**.

**Figure 4:**
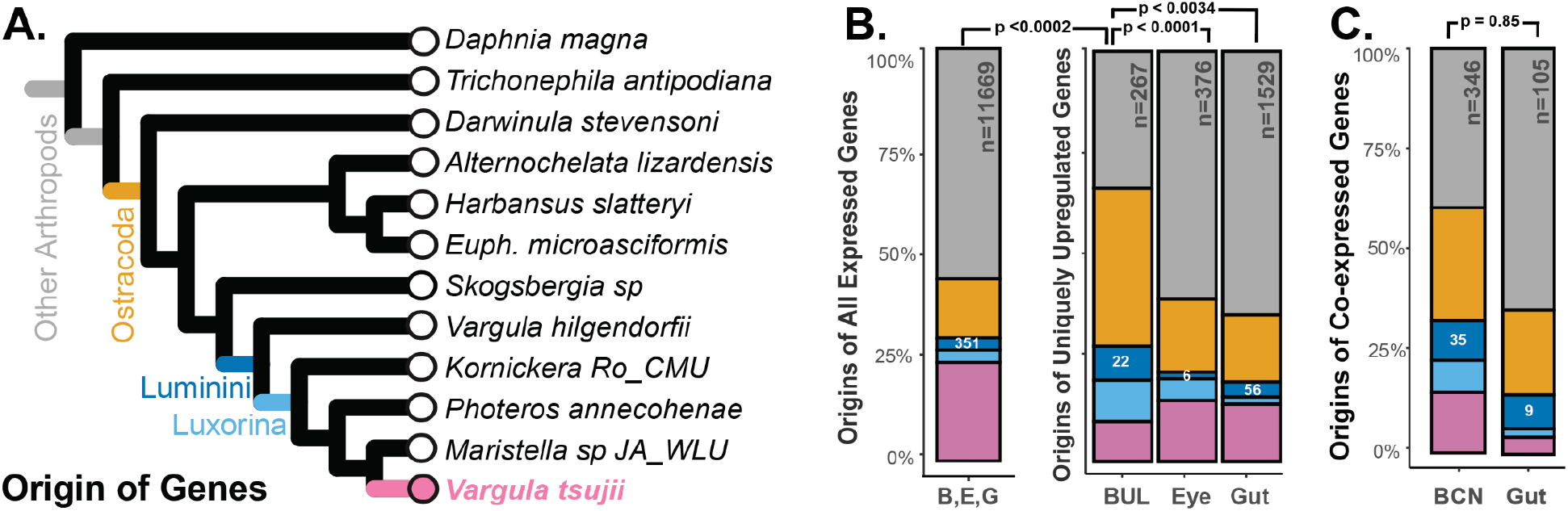
The contribution of novel and conserved genes co-expressed in the BCN and significantly upregulated genes uniquely expressed in the bioluminescent upper lip, compound eye, and gut of *V. tsujii* restricted to *V. tsujii*, Luminini, Luxorina, Ostracoda, and Arthropoda. Conserved ostracoda/arthropoda genes dominate the set of genes in each dataset. **(A)** Phylogeny of bioluminescent and non-bioluminescent ostracods and representatives of other arthropods are used to determine the origin of genes. **(B)** Compared to the complete set of genes differentially expressed in the DGE dataset (minimum of 2 counts or more across biological replicates), only the upregulated genes of the bioluminescent upper lip were characterized by a significantly higher proportion of Luminini-specific genes. Statistically significant pairwise comparison between BUL:DGE Dataset is indicated by p <0.0002. Compared to the proportion of the total number of Luminini genes upregulated in the compound eye and gut, the bioluminescent upper lip had a significantly higher proportion of Luminini genes compared to the gut and compound eye. Statistically significant pairwise comparison between BUL:Eye and BUL:Gut is indicated by p<0.0001 and p<0.0034, respectively. **(C)** Compared to the module expressed in the gut, the BCN contains a higher proportion of Luminini genes indicated, though not statistically significant. *B/BUL - Bioluminescent Upper Lip, E - Eye, G - Gut*.

## Discussion

Through detailed analyses of gene expression, we show that an evolutionarily novel bioluminescence system (22) uses deeply conserved genes of a secretory toolkit to impact ecological interactions by creating light. As such, the secreted bioluminescence of ostracods expands a growing case for the common re-deployment of a conserved secretory toolkit to a new sensory modality, even in secretory innovations across diverse and distantly related taxa. This secretory toolkit has integrated novel genes time and again during evolution to create innovative biological chemistries with dramatic effects on ecological interactions among species.

We found a suite of deeply conserved genes expressed in upper lips and the Bioluminescent Co-Regulatory Network (BCN) to include many toxin-like genes **(Fig. 1A** and **Fig. 3)**. Bioluminescent mucus is conspicuously secreted during predation attempts on Luminini ostracods (35), presumably having impacted predator-prey interactions for hundreds of millions of years (22). However, why fishes spit out ostracods is unknown. Bioluminescence may be aposematic (36), sometimes startle would-be predators (37–39), or makes them more vulnerable to larger predators (‘burglar alarm effect’) (13, 38, 40); but these remain untested hypotheses in ostracods. The mucus produced by ostracods also could contain unpalatable or toxic substances (15), plausible because fish often expel an ostracod during predation attempts (15). We find both a luminous and non-luminous species to express toxin-like genes in their upper lips, with a higher proportion of such genes expressed in the non-light defended *Skogsbergia sp*. **(Fig. 3A and Fig. 3B)**. Notably, toxin-like genes include some related to CAP, which play roles in defense, reproduction, and immune regulation, and include the allergenic Ag5 protein from *Polybia paulista (41)*, (42). Also expressed are CUB domains found in venomous animals (43–45), metallopeptidase M12 genes common in venom (46), and genes from the lipocalin and serine protease families, known for their roles in predation and defense (47). These findings suggest a potential defensive function of ostracod mucus, which should be further investigated through behavioral assays with fish or other predators and with detailed assessments of gene function. Currently, we only have gene expression data for two species, so future work characterizing secretomes across diverse ostracod species could test a hypothesis that the origin of secreted toxins preceded the origin of bioluminescence. Regardless of the outcome of future experiments on function, the expression of these toxin-like genes support a role for long-conserved gene families in secreted bioluminescent systems.

Another class of deeply conserved genes that are highly expressed in the luminous upper lip and BCN are likely involved in mitigating cellular stress, perhaps to tolerate the high secretory load of bioluminescent secretion. Dedicated gland cells that secrete large amounts of proteins or small molecules have a high secretory load compared to other cells and may require a quality control system to accommodate mass production and secretion of products (10, 31, 32). The high demand of light-emitting compounds and other secretory products probably requires activation of cellular and protein stress response mechanisms (ERS response pathways) to increase secretory capacity of cells (10). Having a secretory toolkit to control and regulate cellular processes could facilitate evolutionary transitions from lower-output to high-output secretory systems (8, 10, 31, 32). While we know large volumes of mucus are secreted during ostracod bioluminescence, we know of no similarly high-volume secretory function for non-luminous upper lips, which may function in digestion by smearing mucus on food (20, 48, 49), a far cry from the large, rapid, and voluminous secretions of luminous ostracods during defense or courtship displays (19, 50). Consistent with facilitation of high secretory demand, the bioluminescent upper lip upregulates more secretory toolkit genes than the non-luminous upper lip **(Fig. 3C)**, with several as hub genes, strongly co-expressed with c-luciferase in the BCN **(Fig. 1A)**. The GO enrichment analyses of BCN and luminous upper lip also reflect stress-related processes, such as protein processing and folding, modulation of endoplasmic reticulum stress, apoptosis, redox regulation, and restoring cellular homeostasis; all of which are tightly associated with ER function (**Fig.1A** and ***S1 Appendix, Fig*.*S3*)** (31, 51). Intriguingly, we also found many genes in the BCN implicated in lipid metabolism and transport, which link to protein secretory pathways, mitigate cellular stress, and have been functionally implicated in firefly bioluminescence by providing a source of energy (52, 53). Although the secretory toolkit genes significantly upregulated in the luminous upper lip are expressed in the transcriptome of the non-luminous *Skogsbergia sp*., most of these genes are not significantly upregulated, further supporting the establishment during evolution of a high-output bioluminescent system through the deployment of ancient, conserved machinery for secretory and quality control processes that mitigate cellular stress.

Coupled with legacy elements of a conserved secretory toolkit, we also document the expression of novel genes, which in secretory systems may often enable organisms innovative ways to chemically interact with other organisms and their environment (10, 31, 32). As observed in other secretory and evolutionary novelties (54–57) **(Fig. 4)**, we found many Luminini-specific genes in the BCN and remarkably a high number in the novel bioluminescent upper lip. We also found the interactions of the BCN to be different than co-expression networks of a non-luminous relative, perhaps due to new co-expression patterns in part from novel genes. In addition to c-luciferase, other novel genes could also be functionally critical. For example, about half of the putatively secreted genes upregulated in the bioluminescent upper lip and co-expressed in the BCN are unannotated, with many of these Luminini-specific, suggesting a possible role in bioluminescent phenotypes. Future functional studies aimed at establishing a direct role of novel genes in the evolution and development of cypridinid bioluminescence phenotypes would be valuable.

As cradles of evolutionary innovation, exocrine glands have catalyzed the emergence of diverse biological functions across many unrelated taxa (10). We report gene expression patterns in secreted bioluminescence of ostracods, despite being a lineage-specific innovation, use deeply conserved genes, including toxin-like genes and genes that are involved in secretory processes to protect against cellular stress. These genes may represent an ancient secretory toolkit for high-output secretory cells, similar to gene sets used in the few other secretory innovations studied in sufficient detail (6, 8). Secretory innovations may therefore routinely integrate a similar and conserved secretory toolkit with novel biosynthesis pathways. By leading to the production of myriad new small molecules and proteins, including pheromones, toxins, and bioluminescence, this process of secretory innovation has had a profound impact on ecological interactions throughout evolutionary history.

## Materials and Methods

To understand the contributions of new and existing genes and their co-expression patterns to an evolutionarily novel bioluminescent upper lip gland, we conducted two major classes of analyses. First, we conducted Differential Expression (DE) of upper lips, eyes, and guts of one luminous and one non-luminous species. Second, we quantified gene co-expression networks in the same two species.

### RNA extraction, sequencing, and mapping

For both DE analyses and co-expression analyses, we isolated RNA from animals kept in lab cultures (16) at UCSB. Cultures of luminous *V. tsujii* originated by trapping with bait near Wrigley Marine Lab, Catalina Island, CA (33.444969, -118.484471). Cultures of non-luminous *Skogsbergia sp*. originated by trapping with bait in *Thallasia* beds near Southwater Caye, Belize (16.812833, -88.083154), kept using published methods for *V. tsujii*, except at 27-30C. For Differential Expression experiments, we extracted total RNA using Agencourt RNAClean magnetic beads (Beckman Coulter). For each of the two species, we sampled 5 biological replicates of 3 organs: dissected upper lip, compound eye, and gut of adults; for a total of 30 organ-level expression profiles (2x5x3). We optimized a protocol using RNAClean magnetic beads to extract total RNA (see protocols.io for details). For co-expression analyses, we added 36 additional RNA samples from *V. tsujii*, extracted using TRIzol (Invitrogen). These samples included 12 adult whole bodies, 9 dissected upper lips from adults, and 15 juvenile *V. tsujii* samples comprising three biological replicates from each of five juvenile instar stages (A-I male, A-I female, A-II, A-III, A-IV). All sample information is summarized in (***S1 Appendix, Table S1***). We used 3’ Tag RNA-seq to quantify RNA expression with sequencing at the UT-Austin DNA Core Facility (58). We used published bioinformatics tools (github.com/Eli-Meyer/TagSeq_util) to trim data using BBDUK, remove low-quality reads (<20), and PCR duplicates. We used gmapper to map cleaned reads to a reference transcriptome for *V*.*tsujii* adult whole bodies and juvenile samples, while Bowtie2 was used for the rest of the *V*.*tsujii* and *Skogsbergia sp*. samples (59, 60). For all samples, weak and ambiguous reads were removed if alignments had < 40 matching bp (59).

### Reference transcriptomes, annotation, and gene ontology analyses

For use in both DE analyses and co-expression analyses, we generated and annotated a reference transcriptome for the non-luminous species, *Skogsbergia sp*. Novogene prepared libraries and sequenced libraries using NovaSeq 6000 PE150. We trimmed reads with Trimmomatic (61), assembled with Trinity (v2.1.1), evaluated transcriptome completeness with BUSCO (62) (89.8% using the Arthropoda_odb9 lineage database), then reduced redundancy by merging transcripts > 90% similar using CD-HIT (63). After CD-HIT, we extracted the longest isoform for each gene with Trinotate (v3.2.2) (61) and identified candidate protein-coding regions with TransDecoder (v3.2.2) (64). For the luminous species *V. tsujii*, we used two different transcriptomes, one of which is an existing reference transcriptome (33). We generated Gene Ontology (GO) terms with BLASTX (v2.10.1) with default e-value of 1e-10 and the SwissProt database (downloaded April 2022) (61, 65). We used Trinotate to predict signal peptides, transmembrane domains, and protein domains with signalP (v4.1), TMHMM (v2.0), and Pfam (v35.0), respectively (66) (67) (68). We examined GO enrichment with TopGO (v2.50.0) (69), using a reference transcriptome for the “universe” dataset and various differentially expressed or co-expressed gene sets as the “test” data set. GO terms with singletons were removed. We then reduced the redundancy of GO terms, and clarified GO relationships (e.g. parent-to-child) to one another using the GO-Figure! Python package (70).

### Tissue-level Analyses of Differential Gene Expression

For DE analyses, we determined in each study species the differentially upregulated genes of three tissue types - compound eyes, guts, and upper lips - using five biological replicates for each tissue/species combination (**Dataset S4 A-F, *S1 Appendix, Table S6***). We employed DEseq2 (v1.40.2) on each species separately, assuming a p-value < 0.05 and FC > 1.5 for the significance of differentially expressed genes using the Benjamini-Hochberg method to account for false discovery rate (FDR) (71). Within each species, pairwise comparisons were done across tissue types (i.e., upper lip to compound eye, upper lip to gut, gut to compound eye), but to determine tissue-specific differential upregulation, each tissue was compared to the other two (e.g., upper lip tissue expression was determined by comparing to both compound eye and gut) (***S1 Appendix, Table S6)***. We performed Fisher’s exact tests between species to determine whether there was a statistically significant number of common GO terms between organs (**Dataset S4 J**).

### Constructing Gene Co-Expression Networks

To quantify co-expression, we used WGCNA(v1.72.1) in R to determine weighted co-expression networks in *V. tsujii*, to identify a Bioluminescent Co-regulatory Network (BCN), and to compare the BCN to gene interactions in a non-luminous relative (72). WGCNA facilitates the identification and characterization of co-regulatory networks by clustering genes based on shared co-expression patterns (72). Genes clustered together in the same module (network) are often co-regulated, meaning they are likely involved in similar biological processes (72). For each module, WGCNA identifies ‘hub genes’ - genes with high network connectivity represented by a module membership value (MM > 0.8) - which are considered integral to regulating the expression of other genes in the module and associated with key processes represented by the module (72). The input expression matrix for *V. tsujii* co-regulation included 31,097 genes as rows and 51 samples as columns with samples normalized with the variance-stabilizing transformation in DESeq2. Following the recommendation by WGCNA, genes with zero counts and genes with less than 5 counts in more than 3 samples were removed to reduce noise (72). We checked for batch effects between samples from different runs and extraction methods using DESeq2. Similarities in expression were calculated by Pearson correlation, creating a matrix that was transformed into an adjacency matrix by raising the correlation to a soft threshold power β to preserve the strongest correlations and reduce noise. We chose a soft threshold of 8 using “pickSoftThreshold” and created an adjacency matrix as a “signed” network, where modules correspond to positively correlated genes. This threshold of 8 satisfied the scale-free topology criterion for our data set, where the R^2^ index of scale-free topology is above 0.9 (72). We used a threshold of 0.2 and a minimum module size of 30 to merge similar expression profiles, leading to 13 co-expression modules in *V. tsujii*. Of these, we identified a Bioluminescent Co-Regulatory Network (BCN) as the module containing a c-luciferase shown to function in bioluminescence (26) (**Dataset S1 A**). Construction of co-expression networks for *Skogsbergia sp*. and conservation of BCN orthologs in the networks of the non-luminous relative can be found in (***S1 Appendix, Methods***).

### Determining orthologous genes and comparative transcriptomics

To generate a cross-species expression matrix, we first determined one-to-one orthologs across the reference transcriptomes of *V. tsujii* and *Skogsbergia sp*. using OrthoFinder (73), which infers gene families that originated before each of the ancestral nodes of a species tree. For cross-species comparison, we used only the upper lip, gut, and eye expression data to allow comparable datasets between *V. tsujii* and *Skogsbergia*. To determine the similarity of expression between the upper lips across species, compared to the eye and gut tissue, we performed principal component analysis (PCA) using one-to-one orthologs expressed in both species. To compare across samples, expression counts of a cross-species expression matrix were filtered (expression counts with less than 5 counts in more than 3 samples were removed), and normalized by adding a pseudo count of 1 x 10^−5^ to prevent log_2_(0) scores, followed by a log_2_ transformation using the log2 function in DESeq2 (71). The batch effect caused by multiple species was removed using an empirical Bayes method performed by the ComBat function in sva (74).

### Assessment of clade-specific genes

A recent phylogeny of cypridinid ostracods showed a single origin of bioluminescence and a single subsequent transition to using bioluminescence for courtship (22). To identify genes in *V. tsujii* that originated just before cypridinid bioluminescence (Luminini), just before bioluminescent courtship signaling (Luxorina), or before the origin of ostracods (which we termed ‘Ostracoda’ or ‘All Arthropods’) we used Orthofinder, adding previously published transcriptomes with taxon sampling summarized in (**Dataset S5 A**). We analyzed orthologous groups using KinFin (v1.1) using four user-defined taxon sets: Arthropoda, Ostracoda, Luminini, and Luxorina. Out of the 14690 proteins that clustered into orthogroups, our KinFin analysis identified *V*.*tsujii* proteins restricted to Arthropoda (n = 9182, 62.5 %), Ostracoda (n = 2991, 20.4 %), Luminini (n= 646, 4.4 %), Luxorina (n = 747, 5.8%) and *V*.*tsujii*-specific proteins that were found in orthogroups of two or more sequences (n= 1124, 7.6 %). With the addition of singletons, the number of *V*.*tsujii*-specific proteins totaled a final count of 3905 proteins (22.4 % of *V*.*tsujii* proteome). For *V*.*tsujii*, we determined how many expressed genes are found in each clade and how many are clade-specific genes (without nested genes) (75). We performed Fisher’s exact tests between organs and modules to determine whether there was a statistically significant number of Luminini-specific (without nested proteins) in the BCN and bioluminescent upper lip compared to conserved genes shared with non-luminous ostracods and arthropods (**Dataset S5 B-C**).

### Identifying secretory pathway genes and putative secreted transcripts

To identify secretory pathway genes in the BCN and significantly upregulated in the upper lips of *V*.*tsujii* and *Skogsbergia sp*., we created a reference database of secretory protein pathway genes found in (76). We performed a blast search with an e-value threshold of 1e-5 against this reference database and additionally searched for transcripts with relevant gene annotations and GO terms associated with the secretory protein pathway. To identify putative secreted transcripts, we retrieved all transcripts with a signal peptide and without a transmembrane domain from the set of genes that is both co-expressed with c-luciferase in the BCN and also significantly upregulated in the bioluminescent and non-bioluminescent upper lips. Transcripts were identified as putative toxin-like transcripts if they had a domain present in known toxin protein families related to toxins. We manually searched for InterProIDs for known protein toxin families or domains listed in (7), as summarized in (**Dataset S2 A**) and any GO terms related to toxin and venom (7). Next, to determine if the number of putative toxin-like genes in the bioluminescent upper lip is significant, we compared the number of toxin-like transcripts in the bioluminescent upper lip to the number of toxin-like transcripts found in tissue types not expected to contain toxin-like transcripts (compound eye and gut). We used similar steps to identify putative toxin-like transcripts expressed in the non-bioluminescent upper lip compared to the two organs not predicted to be involved in defense, the compound eye and gut of *Skogsbergia sp*. For each species, we performed Fisher’s exact tests to determine if both upper lips express a significantly higher number of putative toxin-like transcripts compared to the compound eye and gut. We also expanded our search for putative toxin-like transcripts by performing a blast search against the ToxProt database which includes all genes expressed in venomous or poisonous tissues across different phyla, summarized in (**Dataset S2 I**) (77).

## Data Availability

All code, data, figures and tables can be found at https://github.com/lmesrop/BCN_publication. All study data are included in the article and/or supporting information. The data reported in this paper has been deposited in the National Center for Biotechnology Information Short Read Archive, https://www.ncbi.nlm.nih.gov/sra (BioProject: PRJNA1109557)

### Acknowledgements

This study was funded by the Society for the Study of Evolution (SSE) Graduate Research Excellence Grants, Wrigley Institute Graduate Fellowship and the Worster Summer Research Fellowship. Use was made of computational facilities purchased with funds from the National Science Foundation (CNS-1725797) and administered by the Center for Scientific Computing (CSC). The CSC is supported by the California NanoSystems Institute and the Materials Research Science and Engineering Center (MRSEC; NSF DMR 2308708) at UC Santa Barbara. This work was supported by the US National Science Foundation grants DEB-2153773 and IOS-1754770 to THO.

## Supplemental Information

[S1_Appendix](https://docs.google.com/document/d/1S1lNxYX5z1PyWGvV0TNNYvAXs7nqA5KlTA0aMdaFm40/edit?usp=sharing)

