## SupplementalMaterial for "Ancient secretory pathways contributed to the evolutionary origin of an ecologically impactful bioluminescence system"

**This PDF file includes:**

Supporting Methods
Figures S1 to S6
Tables S1
Legends for Datasets S1 to S5
SI References

### Supplemental Material

Additional supplementary material is available online at https://github.com/lmesrop including code, datasets, and figures.

*Investigating the conservation of BCN orthologs in the co-expression networks of the non-luminous relative*

To determine if the BCN shares similarities and is conserved in the co-expression networks of the non-bioluminescent relative, we checked whether genes in the BCN that are orthologous to genes in the *Skogsbergia sp.* transcriptome are conserved in their interactions in co-expression networks of *Skogsbergia sp.* Our rationale is such that if the BCN is conserved in the non-bioluminescent relative, we would expect to find orthologs in one or a few consensus modules. To achieve this, we performed a single network analysis in *Skogsbergia sp.* and determined the number of *Skosbergia sp.* modules the BCN genes are found in. Co-expression networks for *Skogsbergia sp.* were generated with WGCNA using 15 total samples from three tissues (upper lip, gut, and compound eye), each with 5 biological replicates. Following the recommendation by WGCNA, genes with zero counts and genes with less than 5 counts in more than 5 samples were removed to reduce noise (1). Removing genes with zero or low expression led to a matrix of 25,628 genes by 15 samples, which was normalized with the variance-stabilizing transformation function in DESeq2. For *Skogsbergia* we used $\beta$=9 attaining scale-free topology. We used a threshold of 0.3 and a minimum module size of 50 to merge similar expression profiles, leading to 48 co-expression modules for our *Skogsbergia* data. To test a hypothesis of biological significance for co-expression modules, we checked whether modules correlated with eye tissue contain genes related to eye function and development. Next, to determine whether the BCN is conserved or not, we performed a randomization test and asked if 51 BCN orthologs selected at random from all possible one-to-one orthologs expressed are found in fewer modules than a random set of genes. The randomization test was replicated 10,000 times. A custom script was written in R to carry out the randomization test.

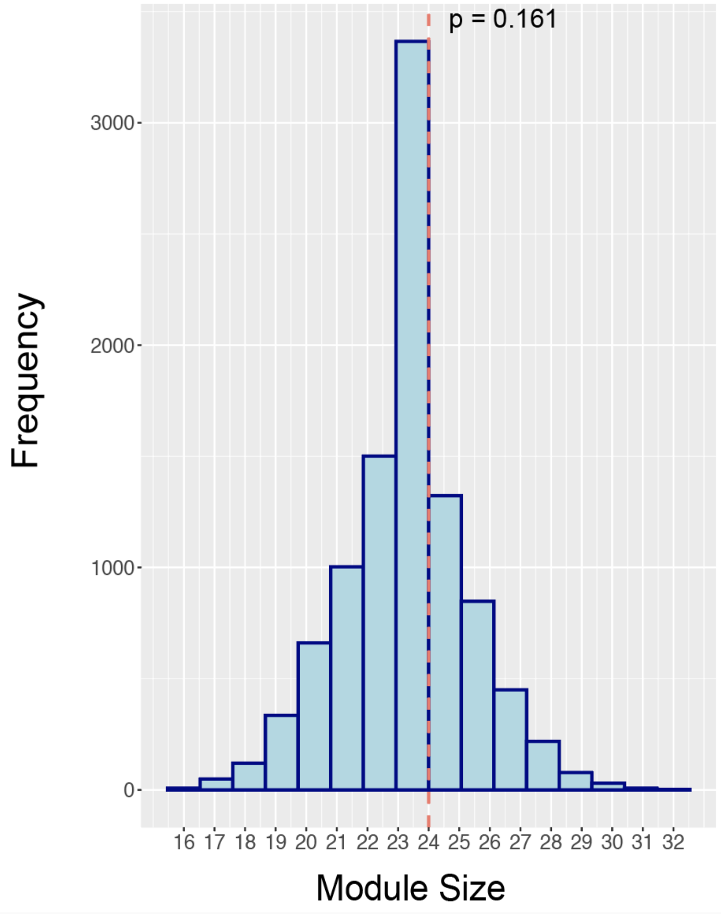


**Figure S1:** Histogram of the results of the randomization test with 10,000 randomizations, examining the distribution of the number of S*kogsbergia sp.* modules (networks) in which the BCN orthologs are found. The rationale is such that if the BCN is conserved in *Skogsbergia sp.* networks we would expect to find orthologs in one or a few consensus modules. We examined the distribution of the number of *Skogsbergia sp.* modules in which one-to-one BCN orthologs are found. The dashed line indicates the experimentally observed number of S*kogsbergia sp.* modules in which one-to-one BCN orthologs are found. The probability of obtaining a distribution in which one-to-one BCN orthologs are found in one or a few consensus modules is not statistically significant ($p = 0.161$).

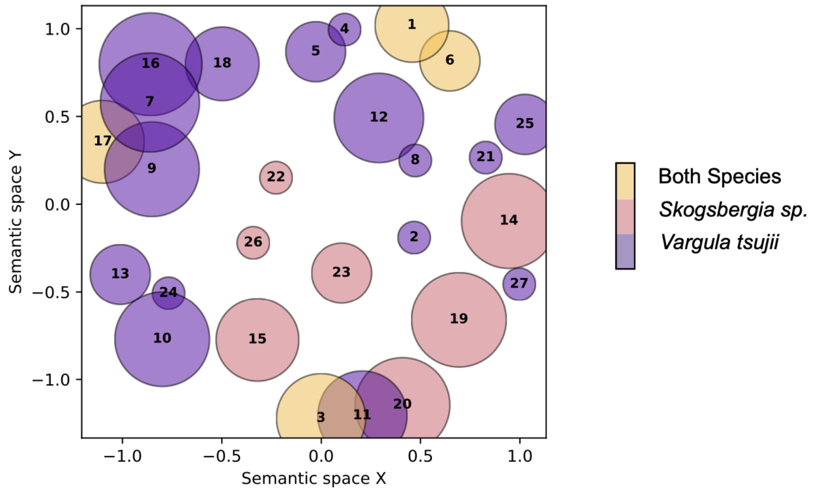


1. proteolysis
2. bioluminescence
3. multicellular organism reproduction
4. peptide amidation
5. chitin metabolic process
6. peptidyl-proline hydroxylation to 4-hydroxy-L-proline
7. positive regulation of cellular pH reduction
8. S-adenosylmethionine biosynthetic process
9. regulation of chloride transport
10. response to fungus
11. digestion
12. calcitriol biosynthetic process from calciol
13. response to cesium ion
14. ion transmembrane transport
15. mating behavior, sex discrimination
16. fibrinolysis
17. regulation of imaginal disc-derived wing size
18. angiotensin-activated signaling pathway
19. stomatogastric nervous system development
20. determination of dorsal identity
21. one-carbon metabolic process
22. epithelial cell proliferation involved in Malpighian tubule morphogenesis
23. cell projection organization
24. cellular response to fluid shear stress
25. collagen catabolic process
26. chemical synaptic transmission
27. oxidation-reduction process

**Figure S2:** Semantic similarity plot of GO biological processes enriched in the bioluminescent upper lip of *Vargula tsujii* and non-bioluminescent upper lip of *Skogsbergia sp.* Only GO terms that are semantically similar across both species are labeled in the plot. Functional enrichment of significantly upregulated was performed separately for each species, GO singletons were removed, and all significant GO terms ($p < 0.05$) were retained in the analysis. Each bubble represents a cluster of similar GO terms summarized by a representative GO term and the color represents the number of GO terms found in each species or both species. Bubble size represents the number of terms in each cluster. GO terms that are similar are plotted closer to one another.

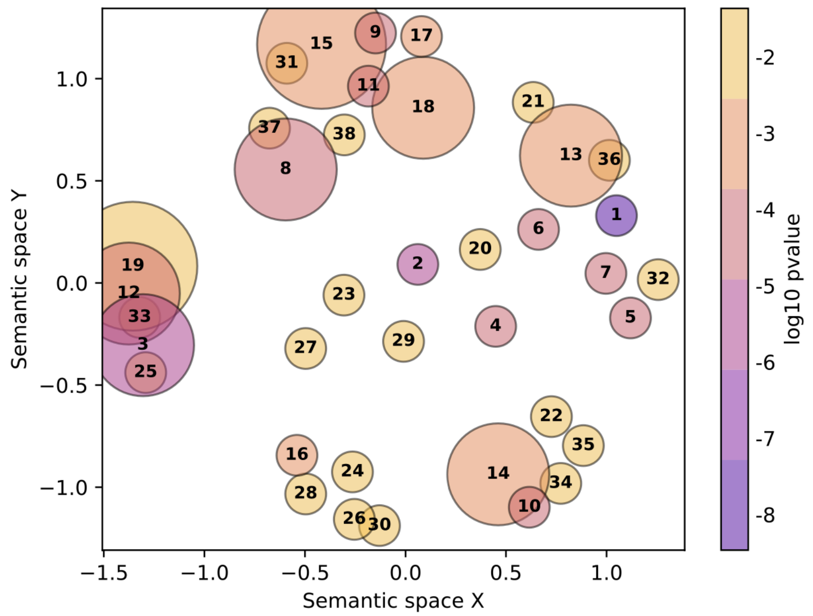


1. proteolysis
2. bioluminescence
3. multicellular organism reproduction
4. peptide amidation
5. chitin metabolic process
6. peptidyl-proline hydroxylation to 4-hydroxy-L-proline
7. S-adenosylmethionine biosynthetic process
8. regulation of chloride transport
9. positive regulation of cellular pH reduction
10. response to fungus
11. positive regulation of synaptic transmission, GABAergic
12. digestion
13. calcitriol biosynthetic process from calciol
14. response to cesium ion
15. fibrinolysis
16. carbon dioxide transport
17. regulation of imaginal disc-derived wing size
18. angiotensin-activated signaling pathway
19. branching involved in ureteric bud morphogenesis
20. one-carbon metabolic process
21. pathogenesis
22. cellular response to fluid shear stress
23. collagen catabolic process
24. bicarbonate transport
25. respiratory gaseous exchange by respiratory system
26. transmembrane transport
27. oxidation-reduction process
28. trehalose transport
29. exogenous drug catabolic process
30. secretion
31. regulation of systemic arterial blood pressure
32. chondroitin sulfate biosynthetic process
33. specification of animal organ identity
34. response to peptide hormone
35. response to pH
36. lipid metabolic process
37. regulation of cilium assembly
38. fibroblast growth factor receptor signaling pathway

**Figure S3:** Semantic similarity plot of GO biological processes enriched in the bioluminescent upper lip of *Vargula tsujii.* Functional enrichment of significantly upregulated was performed separately for each species, GO singletons were removed, and all significant GO terms ($p < 0.05$) were retained in the analysis. Each bubble represents a cluster of similar GO terms summarized by a representative GO term and the color represents the average P-value of the representative GO term. Bubble size represents the number of terms in each cluster. GO terms that are similar are plotted closer to one another.

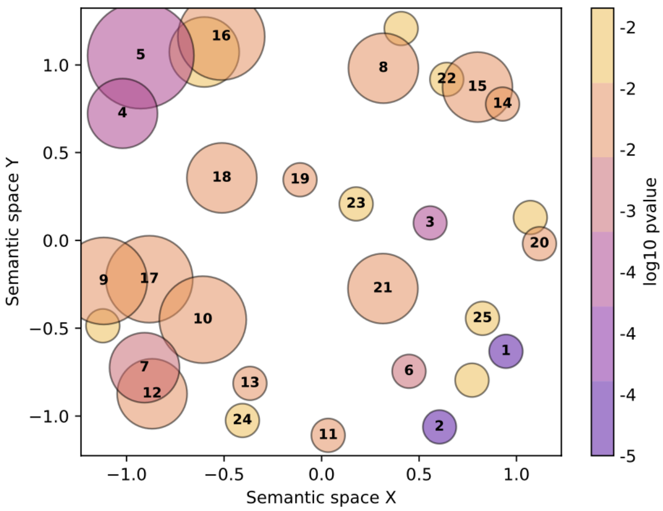


1. peptidyl-proline hydroxylation to 4-hydroxy-L-proline
2. proteolysis
3. ion transmembrane transport
4. mating behavior, sex discrimination
5. memory
6. pantothenate metabolic process
7. negative regulation of tyrosine phosphorylation of STAT protein
8. stomatogastric nervous system development
9. positive regulation of serine-type endopeptidase activity
10. negative regulation of coagulation
11. epidermal growth factor receptor ligand maturation
12. positive regulation of Toll signaling pathway
13. regulation of imaginal disc-derived wing size
14. photoreceptor cell fate determination
15. determination of genital disc primordium
16. determination of dorsal identity
17. regulation of synaptic vesicle endocytosis
18. response to nutrient
19. epithelial cell proliferation involved in Malpighian tubule morphogenesis
20. cell projection organization
21. triglyceride metabolic process
22. compound eye retinal cell programmed cell death
23. chemical synaptic transmission
24. membrane depolarization
25. DNA integration

**Figure S4:** Semantic similarity plot of GO biological processes enriched in the bioluminescent upper lip of *Skogsbergia sp.* Functional enrichment of significantly upregulated was performed separately for each species, GO singletons were removed, and all significant GO terms ($p < 0.05$) were retained in the analysis. Each bubble represents a cluster of similar GO terms summarized by a representative GO term and the color represents the average P-value of the representative GO term. Bubble size represents the number of terms in each cluster. GO terms that are similar are plotted closer to one another.

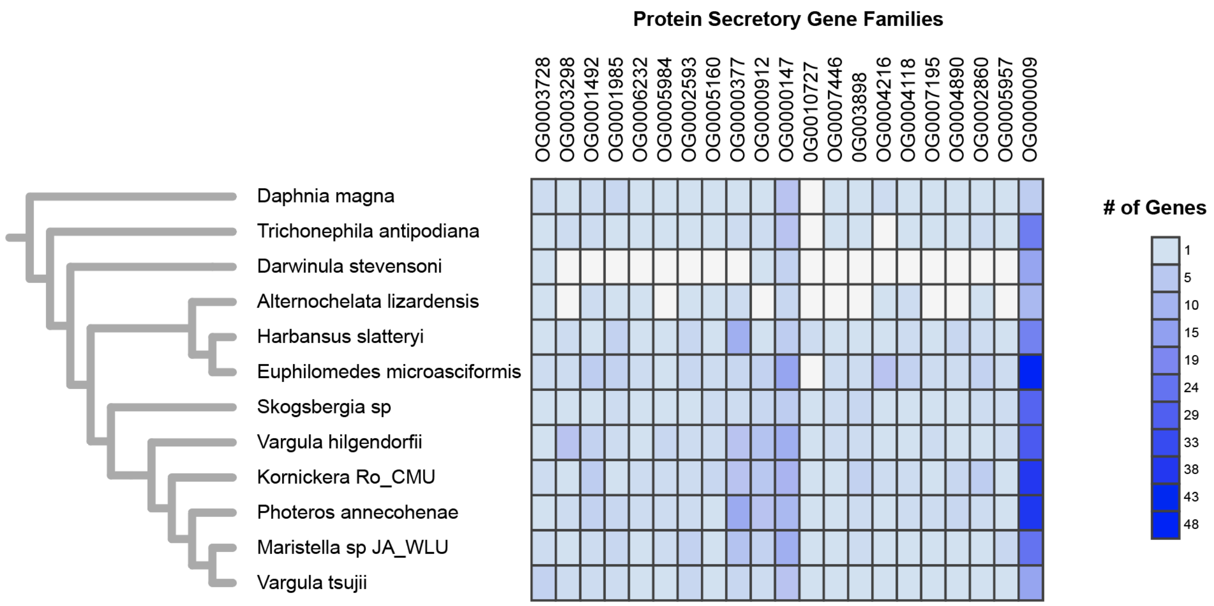


**Figure S5:** The number of genes that belong to known secretory protein gene families significantly upregulated in the bioluminescent upper lip across proteomes of bioluminescent and non-bioluminescent ostracods and other arthropods. White boxes represent the absence of genes.

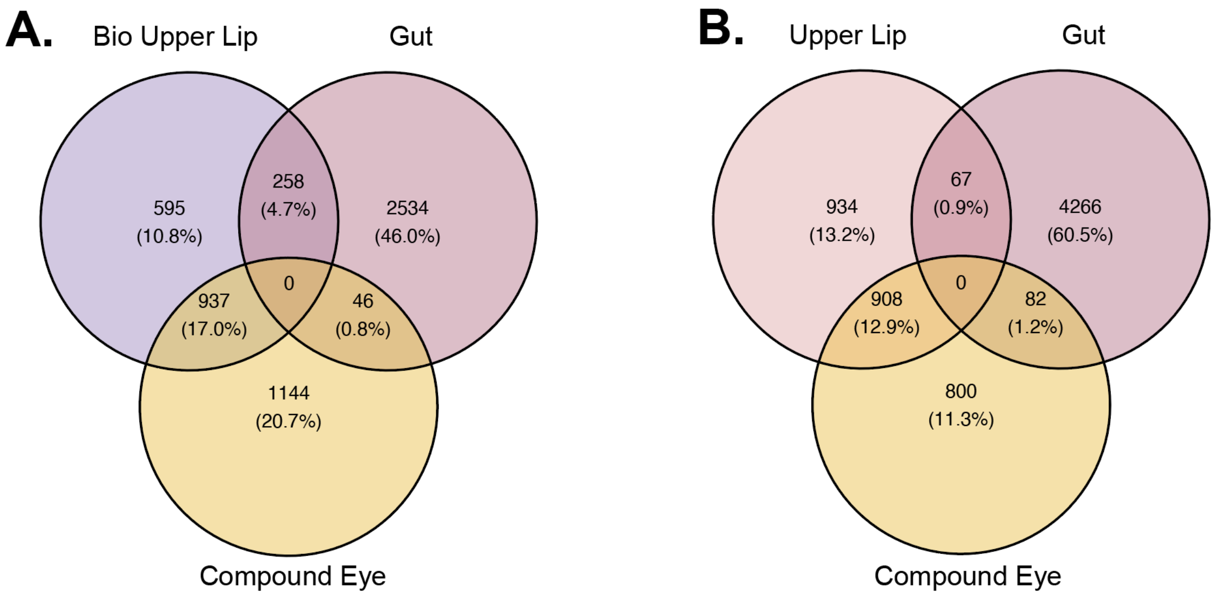


**Figure S6:** Venn diagram of DEG between upper lips, compound eyes, and gut. The numbers in the figure represent the DEG number. **(A)** Venn diagram of DEG between tissue types in the luminous *V.tsujii*. **(B)** Venn diagram of DEG between tissue types in the non-luminous *Skogsbergia sp*.

**Table S1.** Table of all the RNA-seq samples used for both WGCNA and DGE analyses.

| Sample Name | Species | Replicate | Tissue | Description | RNA Method | Analysis Type |
|---|---|---|---|---|---|---|
| Vt.1A.counts.tab | Vargula_tsujii | Vt.1A.counts.tab.rep1 | bio_upper_lip | Adult | Agencourt RNAClean | DE and WGCNA |
| Vt.2A.counts.tab | Vargula_tsujii | Vt.2A.counts.tab.rep2 | bio_upper_lip | Adult | Agencourt RNAClean | DE and WGCNA |
| Vt.3A.counts.tab | Vargula_tsujii | Vt.3A.counts.tab.rep3 | bio_upper_lip | Adult | Agencourt RNAClean | DE and WGCNA |
| Vt.4A.counts.tab | Vargula_tsujii | Vt.4A.counts.tab.rep4 | bio_upper_lip | Adult | Agencourt RNAClean | DE and WGCNA |
| Vt.5A.counts.tab | Vargula_tsujii | Vt.5A.counts.tab.rep5 | bio_upper_lip | Adult | Agencourt RNAClean | DE and WGCNA |
| Vt.1B.counts.tab | Vargula_tsujii | Vt.1B.counts.tab.rep1 | compound eye | Adult | Agencourt RNAClean | DE and WGCNA |
| Vt.2B.counts.tab | Vargula_tsujii | Vt.2B.counts.tab.rep2 | compound eye | Adult | Agencourt RNAClean | DE and WGCNA |
| Vt.3B.counts.tab | Vargula_tsujii | Vt.3B.counts.tab.rep3 | compound eye | Adult | Agencourt RNAClean | DE and WGCNA |
| Vt.4B.counts.tab | Vargula_tsujii | Vt.4B.counts.tab.rep4 | compound eye | Adult | Agencourt RNAClean | DE and WGCNA |
| Vt.5B.counts.tab | Vargula_tsujii | Vt.5B.counts.tab.rep5 | compound eye | Adult | Agencourt RNAClean | DE and WGCNA |

| | | | | | | |
|---|---|---|---|---|---|---|
| Vt.1C.counts.tab | Vargula_tsujii | Vt.1C.counts.tab.rep1 | gut | Adult | Agencourt RNAClean | DE and WGCNA |
| Vt.2C.counts.tab | Vargula_tsujii | Vt.2C.counts.tab.rep2 | gut | Adult | Agencourt RNAClean | DE and WGCNA |
| Vt.3C.counts.tab | Vargula_tsujii | Vt.3C.counts.tab.rep3 | gut | Adult | Agencourt RNAClean | DE and WGCNA |
| Vt.4C.counts.tab | Vargula_tsujii | Vt.4C.counts.tab.rep4 | gut | Adult | Agencourt RNAClean | DE and WGCNA |
| Vt.5C.counts.tab | Vargula_tsujii | Vt.5C.counts.tab.rep5 | gut | Adult | Agencourt RNAClean | DE and WGCNA |
| Sk.10A_fasta97.counts.tab | Skosgbergia_sp | Sk.10A.fasta97.counts.tab.rep5 | upper_lip | Adult | Agencourt RNAClean | DE and WGCNA |
| Sk.6A_fasta97.counts.tab | Skosgbergia_sp | Sk.6A.fasta97.counts.tab.rep1 | upper_lip | Adult | Agencourt RNAClean | DE and WGCNA |
| Sk.7A_fasta97.counts.tab | Skosgbergia_sp | Sk.7A.fasta97.counts.tab.rep2 | upper_lip | Adult | Agencourt RNAClean | DE and WGCNA |
| Sk.8A_fasta97.counts.tab | Skosgbergia_sp | Sk.8A.fasta97.counts.tab.rep3 | upper_lip | Adult | Agencourt RNAClean | DE and WGCNA |
| Sk.9A_fasta97.counts.tab | Skosgbergia_sp | Sk.9A.fasta97.counts.tab.rep4 | upper_lip | Adult | Agencourt RNAClean | DE and WGCNA |
| Sk.10B_fasta97.counts.tab | Skosgbergia_sp | Sk.10B.fasta97.counts.tab.rep5 | compound eye | Adult | Agencourt RNAClean | DE and WGCNA |
| Sk.6B_fasta97.counts.tab | Skosgbergia_sp | Sk.6B.fasta97.counts.tab.rep1 | compound eye | Adult | Agencourt RNAClean | DE and WGCNA |
| Sk.7B_fasta97.counts.tab | Skosgbergia_sp | Sk.7B.fasta97.counts.tab.rep2 | compound eye | Adult | Agencourt RNAClean | DE and WGCNA |

| | | | | | | |
|---|---|---|---|---|---|---|
| Sk.8B_fasta97.counts.tab | Skosgbergia_sp | Sk.8B.fasta97.counts.tab.rep3 | compound eye | Adult | Agencourt RNAClean | DE and WGCNA |
| Sk.9B_fasta97.counts.tab | Skosgbergia_sp | Sk.9B.fasta97.counts.tab.rep4 | compound eye | Adult | Agencourt RNAClean | DE and WGCNA |
| Sk.10C_fasta97.counts.tab | Skosgbergia_sp | Sk.10C.fasta97.counts.tab.rep5 | gut | Adult | Agencourt RNAClean | DE and WGCNA |
| Sk.6C_fasta97.counts.tab | Skosgbergia_sp | Sk.6C.fasta97.counts.tab.rep1 | gut | Adult | Agencourt RNAClean | DE and WGCNA |
| Sk.7C_fasta97.counts.tab | Skosgbergia_sp | Sk.7C.fasta97.counts.tab.rep2 | gut | Adult | Agencourt RNAClean | DE and WGCNA |
| Sk.8C_fasta97.counts.tab | Skosgbergia_sp | Sk.8C.fasta97.counts.tab.rep3 | gut | Adult | Agencourt RNAClean | DE and WGCNA |
| Sk.9C_fasta97.counts.tab | Skosgbergia_sp | Sk.9C.fasta97.counts.tab.rep4 | gut | Adult | Agencourt RNAClean | DE and WGCNA |
| A.1.counts.tab | Vargula_tsujii | A.1.counts.tab.rep1 | whole body | Adult | TRIzol | WGCNA |
| A.2.counts.tab | Vargula_tsujii | A.2.counts.tab.rep2 | whole body | Adult | TRIzol | WGCNA |
| A.3.counts.tab | Vargula_tsujii | A.3.counts.tab.rep3 | whole body | Adult | TRIzol | WGCNA |
| AIF.1.counts.tab | Vargula_tsujii | AIF.1.counts.tab.rep1 | whole body | A-I Instar stage / Female | TRIzol | WGCNA |
| AIF.2.counts.tab | Vargula_tsujii | AIF.2.counts.tab.rep2 | whole body | A-I Instar stage / Female | TRIzol | WGCNA |

| | | | | | | |
|---|---|---|---|---|---|---|
| AIF.3.counts.tab | Vargula_tsujii | AIF.3.counts.tab.rep3 | whole body | A-I Instar stage / Female | TRIzol | WGCNA |
| AII.1.counts.tab | Vargula_tsujii | AII.1.counts.tab.rep1 | whole body | A-II Instar stage | TRIzol | WGCNA |
| AII.2.counts.tab | Vargula_tsujii | AII.2.counts.tab.rep2 | whole body | A-II Instar stage | TRIzol | WGCNA |
| AII.3.counts.tab | Vargula_tsujii | AII.3.counts.tab.rep3 | whole body | A-II Instar stage | TRIzol | WGCNA |
| AIII.1.counts.tab | Vargula_tsujii | AIII.1.counts.tab.rep1 | whole body | A-III Instar stage | TRIzol | WGCNA |
| AIII.2.counts.tab | Vargula_tsujii | AIII.2.counts.tab.rep2 | whole body | A-III Instar stage | TRIzol | WGCNA |
| AIII.3.counts.tab | Vargula_tsujii | AIII.3.counts.tab.rep3 | whole body | A-III Instar stage | TRIzol | WGCNA |
| AIM.1.counts.tab | Vargula_tsujii | AIM.1.counts.tab.rep1 | whole body | A-I Instar stage / Male | TRIzol | WGCNA |
| AIM.2.counts.tab | Vargula_tsujii | AIM.2.counts.tab.rep2 | whole body | A-I Instar stage / Male | TRIzol | WGCNA |
| AIM.3.counts.tab | Vargula_tsujii | AIM.3.counts.tab.rep3 | whole body | A-I Instar stage / Male | TRIzol | WGCNA |
| AIV.1.counts.tab | Vargula_tsujii | AIV.1.counts.tab.rep1 | whole body | A-IV Instar stage | TRIzol | WGCNA |

| | | | | | | |
|---|---|---|---|---|---|---|
| AIV.2.counts.tab | Vargula_tsujii | AIV.2.counts.tab.rep2 | whole body | A-IV Instar stage | TRIzol | WGCNA |
| AIV.3.counts.tab | Vargula_tsujii | AIV.3.counts.tab.rep3 | whole body | A-IV Instar stage | TRIzol | WGCNA |
| C.1.counts.tab | Vargula_tsujii | C.1.counts.tab.rep1 | whole body | Adult Female / Forced bioluminescence | TRIzol | WGCNA |
| C.2.counts.tab | Vargula_tsujii | C.2.counts.tab.rep2 | whole body | Adult Female / Forced bioluminescence | TRIzol | WGCNA |
| C.3.counts.tab | Vargula_tsujii | C.3.counts.tab.rep3 | whole body | Adult Female / Forced bioluminescence | TRIzol | WGCNA |
| E.1.counts.tab | Vargula_tsujii | E.1.counts.tab.rep1 | whole body | Adult Male / Morning | TRIzol | WGCNA |
| E.2.counts.tab | Vargula_tsujii | E.2.counts.tab.rep2 | whole body | Adult Male / Morning | TRIzol | WGCNA |
| E.3.counts.tab | Vargula_tsujii | E.3.counts.tab.rep3 | whole body | Adult Male / Morning | TRIzol | WGCNA |

**Legends for Dataset S1 to S5**

**Dataset S1**
Results of the WGCNA analysis for the BCN. Includes GO enrichment results, genes with high connectivity in the BCN, and tables used for network visualization in Cytoscape.

**Dataset S2**
Table of putative toxin-like transcripts across tissue types for each species.

**Dataset S3**
Table of protein secretory pathway genes across tissue types for each species.

**Dataset S4**
DGE results for the bioluminescent species, *Vargula tsujii,* and non-bioluminescent *Skogsbergia sp.* and also includes the GO enrichment results for the significantly upregulated genes of the bioluminescent upper lip and non-bioluminescent upper lip.

**Dataset S5**
Results from Orthofinder and KinFin analyses which demonstrate the distribution of novel to conserved genes across tissue types and co-expression networks.
